# A Novel Machine Learning Approach Uncovers New and Distinctive Inhibitors for Cyclin-Dependent Kinase 9

**DOI:** 10.1101/2020.03.18.996538

**Authors:** Mariana Assmann, Matthias Bal, Michael Craig, Jarryl D’Oyley, Lawrence Phillips, Hagen Triendl, Paul A. Bates, Usman Bashir, Parminder Ruprah, Noor Shaker, Vid Stojevic

**Affiliations:** GTN LTD, London, UK; Biomolecular Modelling Laboratory, The Francis Crick Institute, London, UK

## Abstract

We present a novel combination of generative and predictive machine learning models for discovering unique protein inhibitors. The new method is assessed on its ability to generate unique inhibitors for the cancer associated protein kinase, CDK9. We validate our method by performing biochemical assays, attaining a hit rate of more than 10%, demonstrating the method to be a notable improvement upon a more standard, and somewhat naive approach. Moreover, we imposed the additional challenge of finding inhibitors that are readily synthesized. Importantly, two new inhibitors are found, with one being distinct from reported CDK9 inhibitors. We discuss the results in the context of modern machine learning principles and the desire expressed by the rational drug design community to secure molecules that are structurally different, yet with high binding affinities, to structurally determined protein targets.

## 1 Introduction

Modern machine learning tools have become impactful, perhaps even essential, in the early stages of the drug design process. Today, deep learning models excel at predicting various chemical properties [1], and deep generative models in conjunction with reinforcement learning are able to efficiently search molecular space for molecules that can optimize several chemical properties simultaneously [2].

The established predictive and generative benchmarks, however, only reflect the real challenges of drug discovery to a limited degree; datasets are usually limited to very small subsets of drug-like molecular space and it is much easier to optimize rule-based or statistical predictors for chemical properties than true values coming from experimental tests. Real-world demonstrations of utility and impact of machine learning tools require far more effort, such as efficient incorporation of appropriate domain knowledge, optimization of computer resources, and, most importantly, the flexibility to address new challenges not reflected in well-curated benchmarks. The purpose of this work is to demonstrate how to address some of the above challenges when pursuing machine-learning driven, budget-constrained, drug discovery programs.

Recently, a machine learning model was employed to find new inhibitors against the kinase target discoidin domain receptor 1, ostensibly demonstrating that such methods can generate new lead candidates in a matter of weeks [3]. However, it was soon pointed out that the newly found inhibitors were very similar to some of the molecules in the training set [4], calling the utility of such methods into question.

In this work we aim to generate new inhibitors for the cyclin-dependent kinase 9 (CDK9). CDK9 is a serine/threonine kinase that was first identified in the early 1990s. It is a member of the CDK family, which plays critical roles in the regulation of cell cycles and transcription. CDK9 is a transcriptional regulator that controls the expression of anti-apoptotic proteins that institute immortality in cancer cells. It interacts with many transcription factors (TFs) and regulates their activities, and it is a recognized molecular target in prostate cancer treatment [5].

We encountered a number of specific challenges in our attempts to find new CDK9 inhibitors:

- Data bias: Publicly available data for binding affinity prediction is very limited, and moreover biased to successes and to non-diversity.
- Binding affinity bias: Most molecules are inactive against a given target, and the chance to randomly find active molecules is very low. A successful model must align with such a prior.
- Generalizability: Machine learning models often work in a certain domain of chemical space but can fail spectacularly outside of that region. This is particularly impactful in activity prediction, where datasets on individual targets are very limited and non-diverse. Generative models can easily generate molecules that are outside the domain to which predictive models generalize. When optimizing for high binding affinity this means that molecules are generated that are falsely predicted to be strongly binding.
- Commercially available molecules: We pursued a cost-efficient strategy using minimal synthesis resources, and exclusively selected molecules from existing molecule libraries. We restricted ourselves to molecules that are in the Enamine Discovery Diversity Set [6], which is a set of molecules already plated ready for assays, and the Enamine REAL database [7], which consists of compounds that can be synthesized in one step. This created a severe restriction on the available chemical search space.

Our first approach was to find new hits against CDK9 by performing virtual screening of the Enamine Discovery Diversity Set using a deep predictive activity model, and simultaneously using predictions against other kinase targets to ensure selectivity in that class. After this approach failed, we analyzed the aforementioned challenges in more detail and developed tools to overcome them. Based on our enhanced approach, we observed, in a second round of synthesis and biochemical testing, a hit rate above 10% (7 hits out of 69 tested molecules), plus active molecules that are clearly distinct from known inhibitors.

## 2 Strategy for finding CDK9 inhibitors

Below we give an abbreviated description of the methodologies developed. More detailed descriptions are provided in Section A.1

### 2.1 A first approach: Virtual screening with a multitask binding affinity model

Our first approach to finding CDK9 inhibitors was based on virtual screening of the Enamine Discovery Diversity library [6]. We used a predictive model that was trained to predict pIC50 values for a set of 14 CDK targets, including CDK9. We then chose molecules that were predicted to be active against CDK9 but not any of the 13 other CDK targets, and had convincing docking poses. Figure 1 shows the pipeline that we used. In this way we chose 207 molecules, given in Section A.8, that were tested in biochemical assay against CDK9. None of the molecules was found to have an inhibition greater than 50% at a concentration of 30 *μ*mol.^1^ In hindsight, we could already identify a problem in our naive machine learning strategy related to optimizing for selectivity. It is well-known that all kinases have very similar functions and, therefore, share a lot of binders. Hence, activity predictions against different CDK targets are strongly correlated, and predictions that a molecule is active against CDK9 but inactive against all other considered targets have a high probability to be wrong on their CDK9 activity prediction. This means that we accidentally picked molecules that have a high uncertainty on their CDK9 activity predictions. Together with the fact that the used datasets over-represent active molecules, this could explain why no inhibitors were found using this approach.

**Figure 1:**
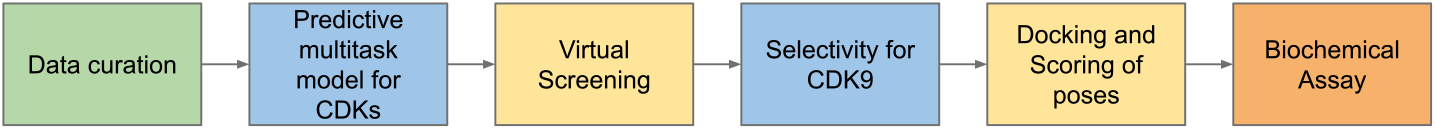
Schematical overview of the pipeline in the first approach. The different parts of the pipeline are indicated by differently colored boxes. Green is for the data curation, blue for predictive training and use of predictive models, yellow indicates filtering processes, and orange is for biochemical testing.

### 2.2 A refined strategy: Optimized inhibitor generation in domain of predictive binding affinity model

#### 2.2.1 Developing a generalizing binding affinity model

One obvious problem with our first attempt was the strong bias toward active components in the dataset. In order to investigate how this bias in the dataset translates into a bias of the predictive model, we decided to investigate the distribution of pIC50 values and compare this to expectation from domain experts to estimate the statistical chance for the model to be accurate. The distribution of expected pIC50 values can be obtained by predicting pIC50 values for randomly sampled molecules from the space of interest (i.e. the Enamine REAL dataset). We found that using a model based on the original dataset roughly three percent of all molecules were classified as active (meaning pIC50 > 7). This is a rate that is orders of magnitude higher than what one would expect in reality, meaning that most molecules that the predictive model identifies as actives must be false positives.

We first tested changing the distribution of predictions by randomly labelling molecules as inactive during training, using either the same molecules or different molecules in each epoch. This naive method unfortunately barely changed the distribution of target values, but created a lot of noise that reduced the performance of the model.

The distribution improved drastically after the negative results for the 207 molecules that had been tested in the first experiment were added to the dataset. The number of active molecules reduced to 0.1 percent, corresponding to an improvement in precision by roughly a factor of 30. Though a ratio of 0.1 percent was still too high, the hope was that it was a sufficient basis for further improvements. This shows that having access to negative results is highly impactful, and suggests an efficient active learning strategy: By synthesizing a few diverse datapoints (in our case a single batch of less than 5% of the dataset size) that the predictive model considers as active, the precision can be greatly improved.

In general we can view this method as an active learning method, meaning a method to acquire new labels in a way that improves the model optimally: If we choose a diverse set of datapoints that are predicted to have high activity and measure their activity against the target, each of those datapoints is either an inhibitor, or, more likely, a datapoint of very high uncertainty whose label improves the predictive model maximally. In contrast, our initial experiments indicated that methods that estimate uncertainty based on a distribution of models are sincerely limited for small and non-diverse datasets. Similarly, typical active learning methods on molecular data seem to give negligible gains compared to random acquisition of datapoints [8]. For activity data our method seems to be much more reliable and leads to drastic model improvements (precision improved by a factor of 30) with less than 5% of newly acquired labels compared to the dataset size.

A second concern was the generalizability of the model. In the first experiment we had simply assumed that the predictive activity model could generalize to the screening dataset. Testing generalizability is already a challenge for small datasets: Scaffold splits, e.g. based on Murcko scaffolds [9] as found in DeepChem [10], do not really capture the problem, as our models score nearly as highly (R2=0.82) as on random splits (R2=0.84).

We addressed this problem by splitting the data manually, using expert knowledge. Since the split was based on the presence of a particular substructure (2-aminopyridine, a common motif in kinase inhibitor design), we refer to it as a substructure split. Inhibitors containing the substructure where assigned to the training set, see Figure 2. This substructure split allow us to evaluate the ability of the model to predict against new scaffolds, and in contrast to automated Murcko scaffold splits the advantage here is that training and test set are consistently separated by certain motives. The substructure split proved to be a much more exacting test of generalizability: validation R2 was 0.32 for our predictive model.

**Figure 2:**
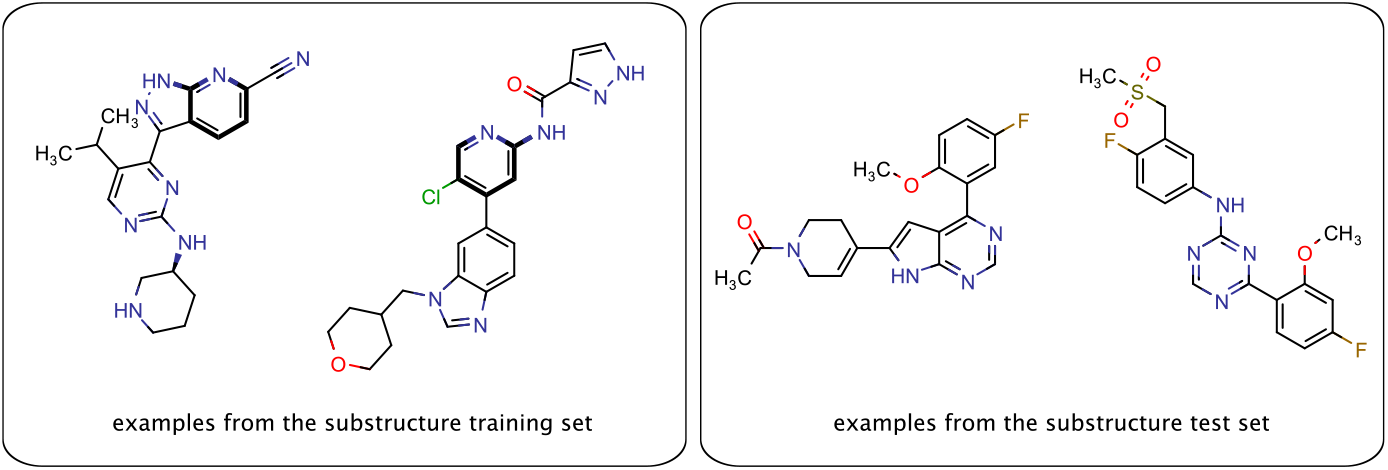
Some examples from the CDK9 activity dataset and their structures. The two molecules from the training set contain the 2-aminopyridine substructure (highlighted in bold) and the test set contains molecules without this moiety.

In order to improve the generalizability of the model, we used Deep Graph Infomax style unsupervised pre-training [11, 12, 13] regularized by early stopping. With this method, we could raise the performance to R2=0.48 on the substructure split, while performance on random splits only slightly decayed (from R2=0.84 to R2=0.78).

#### 2.2.2 Guided molecule generation

One of the major limitations of the first experiment was its reliance on virtually screening a specific dataset. A guided generative model brings more flexibility and possibilities to find new inhibitors. A generative model learns the distribution of a specific dataset by training to produce data similar to samples from the dataset. In our use case the objective of this pre-training is to learn to produce valid and drug-like molecules with a good chance of being CDK9 inhibitors. In a second training sequence the distribution represented by the generative model gets optimized with respect to certain their inhibition qualities. Both training steps are discussed in more detail in Section A.2.

In order to produce a high ratio of molecules that the predictive model can generalize to, we chose the Enamine kinase library [14] and added the molecules from the CDK9 binding affinity dataset as a promising starting point for this objective. For consistency between the generative and predictive model, we also used the Enamine kinase library and the binding affinity dataset together for the pre-training of the predictive model.

Since even in the optimistic case the precision of the predictive model is rather low, the generative model might optimize for false positives, as these might be more abundant than true positives. Therefore it is important to estimate the reliability of binding affinity prediction on generated molecules. Our first attempts using standard uncertainty measures like training a Bayesian neural network or using Gaussian Processes on the features extracted for the predictive model did not lead to solid results, mostly because the dataset was too small and non-diverse to find a reliable probability distribution.

It was noticed before that out-of-distribution data points are a major source of uncertainty [15]. If generated molecules are outside the training set distribution, binding affinity prediction will be unreliable. We therefore developed an out-of-distribution classifier consisting of a support vector machine on top of the final-layer features of the predictive model. We trained this model to distinguish molecules from the CDK9 training set against generic molecules from the REAL dataset. The classifier achieved a performance of 0.98 for both precision and recall. We assumed that the predictive model should generalize to generated molecules classified as *in-distribution* with the CDK9 training set, and therefore used the classifier as an additional filter.

#### 2.2.3 Finding candidate CDK9 inhibitors

For the refined strategy we updated the CDK9 data by adding the inactive compounds from the first approach. With this dataset we trained an ensemble of predictive models that was used to provide the generative model with feedback during training.

In order to keep synthesis cheap we identified for each generated molecule nearest neighbors in the REAL database and repeated the ensemble analysis for those compounds. They were scored with the aforementioned ensemble of predicted models, and filtered by physico-chemical properties and out-of-distribution classification. From the remaining compounds, 69 were manually selected for testing in biochemical assays. An overview of the procedure can be found in Fig. 3, and the details for the whole procedure are outlined in Section A.7.

**Figure 3:**
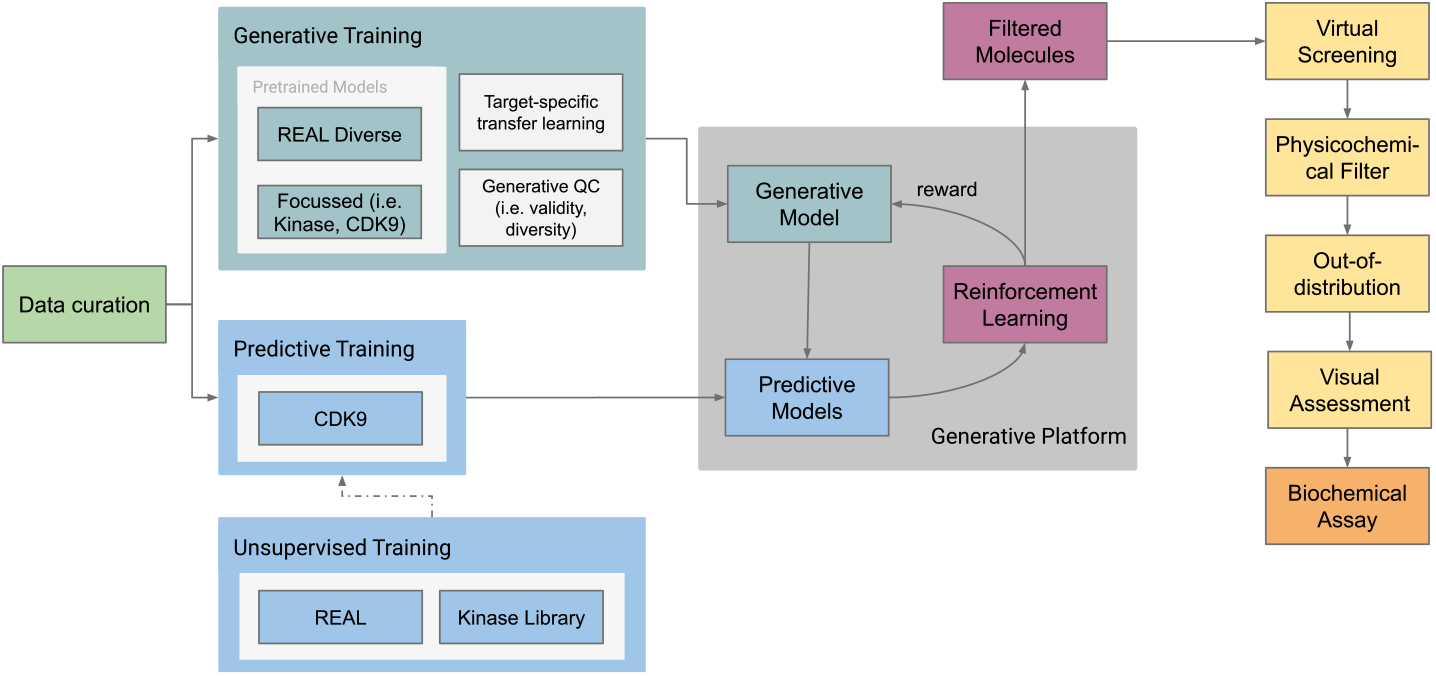
Overview of the round 2 procedure. We use the same colour scheme as in Figure 1 with the addition of teal indicating generative training and models, and magenta for reinforcement learning.

From the 69 tested compounds, seven showed significant activity. These are depicted in Figure 4, and their corresponding activity data can be found in Table 1.

**Table 1:** Properties of the molecules with significant activity shown in Figure 4. % Activity is the percentage of protein activity in the presence of the inhibitor. In the last column we report the highest Tanimoto similarity of the molecule that was found by comparing to all molecules in the CDK9 binding affinity dataset.

| Molecule | % Activity<br>at 10μM/30μM | Highest<br>similarity |
| --- | --- | --- |
| 1 | 25 / 6.4 | 0.59 |
| 2 | 34 / 12 | 0.51 |
| 3 | 55 / 16 | 0.52 |
| 4 | 55 / 21 | 0.47 |
| 5 | 63 / 37 | 0.45 |
| 6 | 64 / 38 | 0.48 |
| 7 | 61 / 44 | 0.45 |

**Figure 4:**
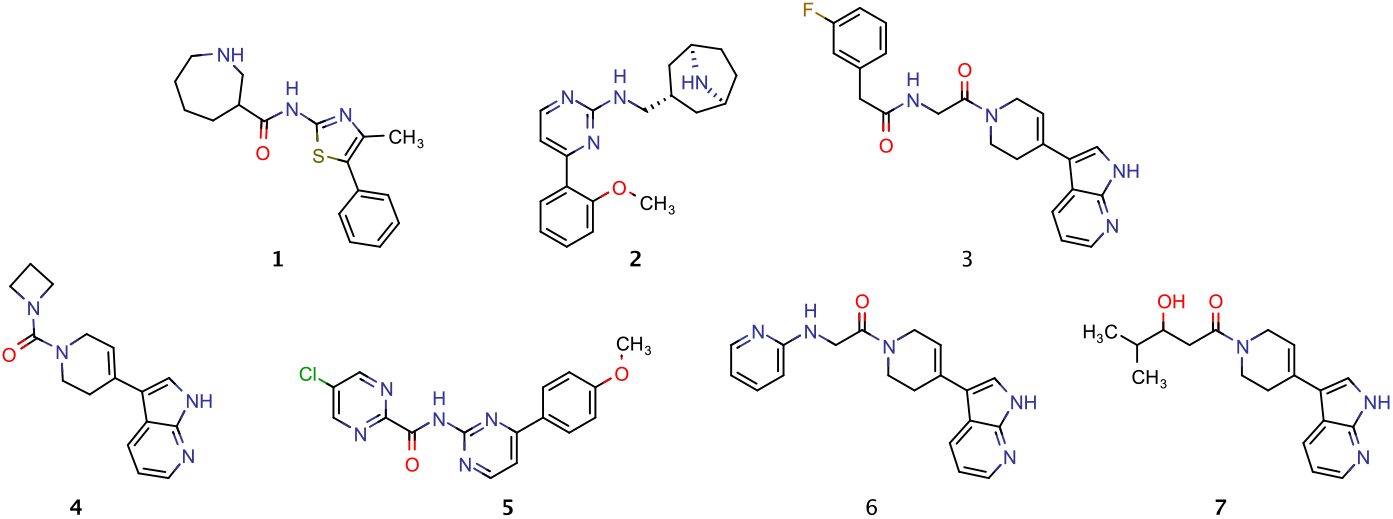
Structures of the molecules that showed significant activity.

#### 2.2.4 Novelty of hits

All seven active molecules have a maximum Tanimoto similarity of 0.6 to any molecule in the CDK9 binding affinity training set, see Table 1. This shows that there is a significant difference in the structures.

More generally, the novelty of hits was investigated with PubChem [16] (accessed September 2019) which we used to first find compounds similar to the seven active molecules. None of the molecules were in PubChem, that is, there was no hit with Tanimoto similarity of 1. We therefore identified all compounds with a Tanimoto similarity greater than 0.9 and checked if the obtained compounds can be found in any patent that covers the CDK family.

For our most tightly bound compound, there were 184 molecules with a Tanimoto similarity > 0.9. Of these compounds only 2 were registered in a patent (reported as D2 dopinergic receptor agonists, US5068325) with a chemotype distinct from molecule 1.

For molecule 2, there were 584 molecules with a Tanimoto similarity > 0.9. Of these compounds 88 were registered in patents. Two patents with compounds with a Tanimoto similarity > 0.94 were analyzed further. The molecules in both patents (US2015038516 and US2009221581) have examples of inhibitors of CDK9. There is very little difference between 2 and the Markush structure in the patent and therefore it is unsurprising the 2 is a CDK9 inhibitor. The examples in these patents are not in the GoStar dataset [17], from which we got our data, and were not part of the CDK9 dataset used for training.

The molecules similar to the other five active molecules can be found in patents covering other proteins than CDKs, such as Inducible I kappa-B kinase (IKK-e), TANK-binding kinase 1 (TBK1), Kallikrein-1 (KLK1), Rho-associated coiled-coil-containing protein kinase 1 (ROCK1). In cases where the similarity searches yielded more than 100 hits, we looked at patents of the first 50 compounds with the closest similarities.

Most importantly, in molecule 1 we have identified a double-digit micromolar molecule with clear patent space as a CDK9 inhibitor that would serve as promising ‘hit’ for further optimization.

## 3 Discussion

In this work, we showed how a novel combination of machine learning models can be employed to generate new kinase inhibitors, and found a hit rate of more than 10% (7 out of 69 compounds) as tested on CDK9. Moreover, we believe our new machine learning method to have generic properties, and therefore, is likely to be equally productive on other protein target classes for which experimentally determined protein structures are available. Prior to our current machine learning model, a simpler method was unsuccessful and led to 207 molecules tested in biochemical assays to be inactive. Even if we count the first, unsuccessful attempt, the hit rate is still acceptable (> 2.5%). However, we are confident that further rounds of testing would keep a hit rate similar to 10%, as we only improve the data with each round.

All of the seven hits are distinct from our CDK9 activity dataset, and only one of them turned out to be a known CDK9 inhibitor. These results were achieved while constraining synthesis to very easily synthesizable compounds from the REAL database. Now that we have validated our method, we expect much better performance in the absence of such restrictions using bespoke synthesis.

During this work, we identified several challenges that needed to be addressed to be successful: First, the bias in activity datasets requires active learning to give a solid basis for a predictive activity model. We found that synthesizing cheap molecules that are suspected by the predictive model to be active gives the best ‘active learning’ results to improve the model. We also found that it is challenging to create a test set that captures the challenges of generalizing to larger portions of chemical space, and we used extensive expert knowledge to do so. In our experience, no available benchmark dataset or automated splitting method could create a challenge of similar quality or difficulty. Perhaps most importantly, using several different machine learning models together poses additional challenges, as the generative model can easily optimize for molecules that the predictive model is very uncertain about. Unsupervised pre-training gave us more robust representations of molecules, while out-of-distribution classification allowed to avoid regions of high uncertainty.

Our approach has several limitations that will prove even more challenging to overcome. Our machine-learning-based approach could only be successful with the amount of data available, which even for the CDK9 target tested here proved to be challenging. On less explored targets machine learning has to be combined with domain knowledge and computational methods in a more creative manner. This is inevitable when exploring first-in-class targets.

Though we found new inhibitors for CDK9, kinase inhibitors are infamously promiscuous, i.e. often bind to various other kinases, which leads to toxicity issues in later drug discovery stages. We were unable to solve the major challenge of producing selective inhibitors given the amount of data available. We believe that this poses a sincere restriction in machine-learning-based drug design and needs to be addressed either with substantially larger datasets or using domain-knowledge-based methods embedded more purposefully in the machine learning method.

## Supporting information

CDK9 activity dataset

## Acknowledgments

We would like to thank Jarvist Moore Frost for useful discussions and feedback.

## Author contributions

The predictive model and the unsupervised pre-training was developed jointly by M.B., L.P. and H.T.. M.A. curated the data, performed the training of the models and the post-processing of the generated compounds. M.B. suggested and developed the unsupervised learning approach for the predictive model. M.C. developed the generative and reinforcement learning algorithms, and performed the training of the models. J.D. created the split to test generalizability of the predictive model, developed the filters for the generated compounds and selected the compounds for testing. L.P. suggested and developed the out-of-distribution classifier and developed the hyperparameter tuning platform for the models. H.T. led the critical analysis of the first approach and designed the strategy for the second approach. He suggested and performed the bias analysis for the predictive model. H.T., M.A. and J.D. wrote the manuscript. P.A.B. advised on the general strategy and aided in editing of the manuscript. U.B., P.R., N.S. and V.S. supervised the work.

## A Supplementary Material

## A.1 Predictive binding affinity model

Molecules can be conveniently encoded as SMILES strings (Simplified molecular-input line-entry system) or as molecular graphs. We use molecular graphs dressed with additional atomic features as input to the predictive model. We then train a graph-convolutional neural network, cf. [18] and references therein, with an architecture similar to Ref. [19].

## A.1.1 Data curation

All activity data was obtained from the GoStar database [17], which predominately contains patent data along with some datapoints from published articles. The data consists of molecules represented by SMILES strings with one measurement against a target protein per data point, the corresponding reference and specifics about the assay. The data cleaning and filtering process was guided by Refs. [20, 21], and adapted to the GoStar data with the following steps:

- Unifying the protein names to one naming system
- Canonicalizing SMILES using the RDKit [22], converting salts to the parent SMILES, and removing entries with invalid SMILES.
- Standardizing activities and units, and removing datapoints that don’t have concentration measurements.
- Removing impossible activity values (e.g. negative pIC50 values) which were found to be errors in copying the data from the original patents, extremely high or low values (such as pIC50 > 10, pIC50 < 2), and unclear values (e.g. when only an upper or lower bound was given).
- Removing all datapoints for which not a single activity measure was given but a range in which the activity was found to be in.
- Removing duplicates with the same SMILES, activity value and reference.
- If there was a duplicated value but from different publications, remove the newer one assuming that this is very likely a citation of the older publication.
- Any remaining duplicated SMILES were manually checked, the corresponding activity values were averaged or removed.

For the multitask dataset in our first approach, we started with a set of 38 cyclin-dependent kinases, that is all the cyclin-dependent kinases that were available in GoStar and had at least 20 usable measurements after the filter process. This resulted in a sparse dataset of 59,375 compounds that have a measurement against at least one kinase. In order to reduce sparsity, the dataset was reduced to 19,723 compounds by filtering out the kinases that had less than 500 measurements against them. The kinases that were left are: CDK1, CDK2, CDK2-CyclinA, CDK2-CyclinB, CDK3-CyclinE1, CDK4, CDK4-CyclinD, CDK4-CyclinD1, CDK5, CDK6-CyclinD2, CDK6-CyclinD3, CDK9, CDK9-CyclinT1.

For the CDK9 dataset, we used the measurements for CDK9 and CDK9-CyclinT1 from the multitask dataset, and added updated data from the GoStar database. After cleaning and filtering, the dataset comprised 4309 compounds. To this data, we added the compounds that were found to be negative in our first round of testing. Since the exact pIC50 values for these inactive compounds is unknown, all of them were given the same activity values. In order to choose that value, we tried different pIC50 values between 1 and 4, since any pIC50 < 5 indicates an inactive molecule. We found that in that range the value does not influence the distribution of larger pIC50 value predictions significantly, and, therefore, also not the chance of a molecule being predicted active. We choose to assign a pIC50 of 4 to all tested inactive compounds.

## A.1.2 Featurization

The SMILES strings in the dataset are converted with the help of the RDKit into molecular graphs of atoms connected by bonds. With the RDKit we compute the following atomic features: atom type, number of bonds to other atoms, number of hydrogen atoms bound to the atom, type of hybridization, formal charge and whether the atom is aromatic. For the edges we used only one feature, the bond type. Non-binary categorical features where cast to one-hot vectors.

## A.1.3 Model Architecture

We input the graphs in sparse representation using node (***a***) and edge (***e***) feature vectors

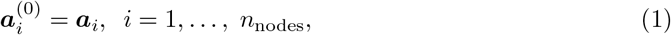

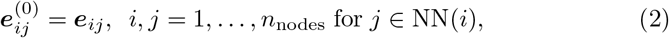

where j belongs to the set of nearest-neighbors NN(i) of i. For each chemical graph, we encode the atom-based features in ***a***_*i*_. The bond type is one-hot encoded in ***e***_*ij*_. Graph-convolutional models are based on a message-passing framework and consist of alternating message-passing steps to process local information, and (optionally) pooling steps to reduce the graph to a simpler sub-graph. A read-out phase gathers the node features, and computes a feature vector of the entire graph that is fed to a perceptron layer for the final prediction step.

In this work we use pair-message and dual-message graph-convolutional layers as discussed in Ref. [18, 19], which are capable of taking both node and edge features into account. The layer computes an aggregate message 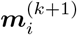 from all neighboring source nodes *j* ∈ NN(*i*) to a target node using a fully-connected neural network *f*_**W**_ acting on the source node features 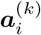 and the edge features 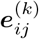 of the connecting edge. To the aggregated result, a self-message from the original node features

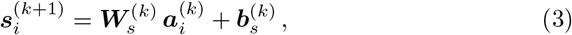

with ***W***_*s*_ being the weights and ***b***_*s*_ being the bias in the linear layer, was added. New node features are computed by applying batch norm (BN) and a ReLU non-linearity, i.e.

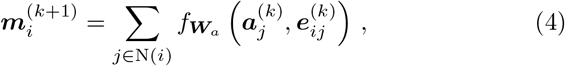

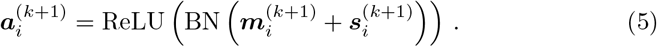

In pair-message graph-convolutional layers the edge features are not updated and remain constant during message-passing. In the dual-message graph-convolutional layers, edge features are also updated with the feature vectors of nodes at the end points of the edge via

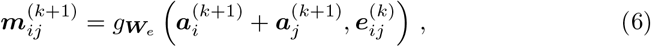

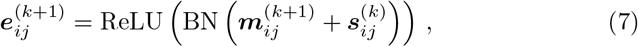

where *g* is a fully-connected neural network and 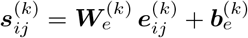 is the edge feature self-message.

In our initial tests both pair-message and dual-message networks performed comparably on the task of pIC50 prediction. We therefore settled for the more economic pair-message network for most predictive models, but used a dual-message network in the ensemble approach.

## A.1.4 Hyperparameter tuning and measuring generalizability

In all of our models we tuned relevant hyperparameters using the hyperband algorithm [23]. Performance of the model was measured in terms of the R2 metric

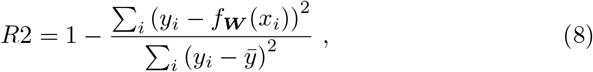

where *f_W_* is the predictive model. The optimal parameters we found for our models are given in Table S1.

## A.1.5 Pre-training

Part of the challenge of generalizing to new data is that the predictive model has never actually learned a useful representation for such data. Therefore it can help to separate learning a meaningful representation of the data before training on the actual prediction task. We choose for this pre-training step an unsupervised learning task, i.e. a task that does not require labels, so that we can freely choose the training set for this first training step. As unsupervised pre-training task we choose mutual information maximization between the neural network layers and the output layer, as explored in Refs. [11, 12, 13], using the negative of the Jensen-Shannon approximation of the mutual information as loss function^2^

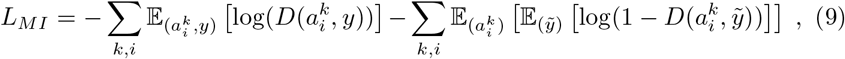

where *D* is the discriminator that attempts to discriminate whether the activation 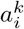 at node *i* and layer *k* and the final layer representation *y* belong to the same data point. We choose as discriminator a simple bilinear layer

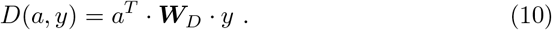

We tuned the hyperparameters of pre-training based on the performance on the downstream task of activity prediction. We observed a high variability between different pre-training runs that led to statistically significant difference in that performance. Therefore we pre-trained a batch of models and then decided on the best pre-trained model using the best performance evaluated by training five different models on the downstream task for each pre-trained model. We also observed that early stopping was crucial in order for the pre-training to have a positive impact. We determined the early stopping parameter together with the other hyperparameters. Using these techniques we could improve performance with the substructure split by 50% (from R2=0.32 to R2=0.48), while performance on random splits slightly declined (from R2=0.84 to R2=0.78).

## A.2 Generative model

Our generative model is an Recurrent Neural Network (RNN) with stacked Long-Short-Term Memory (LSTM) cells [24] that in a pre-training step learns a prior distribution over molecules by maximum likelihood estimation, minimizing

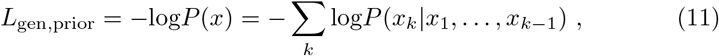

with stochastic gradient descent.

We subsequently optimize the generative model to produce active molecules using the reinforcement learning method Reinvent [25]: The network learns a probability distribution whose likelihood log*P*(*x*) differs from the previously learned prior likelihood log*P*_prior_(*x*) by a term proportional to the scoring function *S*(*x*), using the standard mean-squared error (MSE) loss

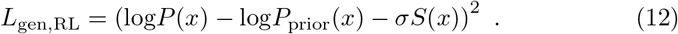

The scoring function *S*(*x*) is in our case determined by the activity of the molecule *x* that is predicted by the predictive model trained prior to the generation step. *S*(*x*) is one if the molecule is predicted to be active (pIC50≥7) and otherwise zero.

## A.3 Out-of-distribution classifier

In order to avoid the generative model to create adversarial examples for the predictive model that score highly but are distinct from the CDK9 dataset, we used an out-of-distribution classifier to classify whether molecules are similar to the CDK9 dataset or not. We used this classifier as an additional filter in selecting candidate CDK9 inhibitors.

The out-of-distribution classifier is a standard support vector machine (SVM) with a linear kernel on top of the final-layer features of the activity predictive model. The molecules in the CDK9 activity dataset served as data labeled *in distribution*. For *out of distribution* data we randomly sampled an equal amount of data from Enamine’s REAL library. The dataset we created turned out to be essentially linearly separable, and the support vector machine achieved an accuracy of 0.98, with an equal percentage of false positives and false negatives.

**Table S1:** Graph-convolutional model hyperparameters of the predictive models used in this work.

| Model | Unsupervised Data | Training Data | Node Channels | Edge Channels | NN Channels |
| --- | --- | --- | --- | --- | --- |
| **First approach** |  |  |  |  |  |
| *Multitask* |  | multitask | 512 |  | [512, 512] |
| **Second approach** |  |  |  |  |  |
| *Supervised* |  | CDK9 | [512, 512] | [256, 256] | [256, 256] |
| *Unsupervised1* | REAL | CDK9 | [512, 512, 512] |  | [128, 128] |
| *Unsupervised2* | First 6 RL Runs | CDK9 | [512, 512] |  | [128, 128] |
| *Unsupervised3* | All RL Runs | CDK9 | [512, 512, 512] |  | [128, 128] |

## A.4 Hyperparameters used in this work

## A.5 Docking details

We performed docking of the molecules found in our first approach into one selected CDK9 protein to be able to visually analyze the binding quality. This was used together with the predicted pIC50 values to select the molecules to be tested experimentally. We used the RDKit to calculate conformations from the SMILES representation of the molecules. The obtained conformations were protonated with MoKa [26], and docked using FLAPdock [27, 28]. As the protein receptor for docking, we used 4BCJ, obtained from the PDBbind database [29, 30, 31]. The protein was prepared for docking using pdbfixer [32]. The pocket was chosen to be of a size of 4 Å around the crystal ligand. As parameters for docking we chose to optimize the docked ligands and to get the best quality of the poses. We chose the “CRY” probe instead of “DRY”, and used the “ampq” keyword to include energetic terms. The eight best poses were calculated.

## A.6 Details of virtual screening approach

Here we give the details of the our first, virtual screening approach with a multitask pIC50 model. A graphical overview of the approach is shown in Figure 1.

As predictive model we used a multitask model that learned pIC50 predictions for a set of 14 cyclin-dependent kinases, including CDK9 and CDK9-Cyclin-T1 (for details on the multitask dataset see Section A.1.1). After hyperparameter optimization, we obtained a model with mean absolute error of 0.46 for CDK9 and 0.43 for CDK9-CyclinT1. Screening the Enamine Discovery Diversity library (50k compounds) across the CDK family yielded around 600 molecules that were predicted to have a pIC50 against CDK9 greater than seven. Those 600 molecules were filtered based on highest selectivity for CDK9 over the rest of the CDK family. Further analysis was done by docking into the 4BCJ protein. The details for the docking procedure can be found in Section A.5. The docked poses were additionally scored with Shape-It [33], and protein-ligand interaction fingerprints (PLIFs) similarities with the 4BCJ x-ray crystal ligand as a reference. The PLIFs were obtained from the output of the protein–ligand interaction profiler (PLIP, Ref. [34]), and Tanimoto similarity to the PLIF of the reference molecules were calculated. For each compound, a visual analysis of the eight best docking poses was done. The compounds were filtered considering the calculated scores, and by manually determining the quality of the docking pose. 207 compounds, listed in Section A.8, where selected and tested in biochemical assays against CDK9. None of the molecules was found to have an inhibition greater than 50% at a concentration of 30 *μ*mol.

## A.7 Details of selection procedure of candidate CDK9 inhibitors

This section outlines the details of our refined strategy; a graphical overview over the whole refined procedure pipeline can be found in Figure 3.

The dataset for this second approach contains 4309 compounds from literature tested against CDK9 and CDK9-CyclinT1. To this, the 207 inactive compounds from the earlier test round where added. We call this the CDK9 dataset. This dataset was used to train predictive, graph-convolutional models using random splits.

We used an ensemble of predictive models, both an ensemble of three models to determine the reward in the generative loop and an ensemble of seven predictive models to select promising candidate inhibitors. The ensemble to determine the reward in the reinforcement learning (RL) algorithm consisted of one model just trained on the CDK9 dataset, and two models pre-trained using unsupervised learning and then fine-tuned on different random splits.

To allow for more diversity in the generative run, we used models from 15 different iterations of the pre-training for the reinforcement learning. With those models, we created 39 RL runs that also used different number of steps, that is 1000, 2000, or 5000. The reward in all RL trainings was given by scoring the generated molecules with the three predictive models as explained above, where a reward was given if one out of the three models predicted a pIC50 greater than 7. The molecules from all the runs were collected and duplicates were removed, yielding a total of 4422 novel molecules.

We used all generated compounds to search for their nearest neighbors in REAL using the KNIME [35] node for Chemspace, and from each hit picked the 10 nearest neighbors with a Tanimoto similarity greater than 0.7. This yielded approx. 17,000 molecules. From those, duplicates were removed, and molecules were filtered for physico-chemical properties calculated with the RDKit. We selected molecules that have molecular weight greater than 250, a total polar surface area between 70 and 220, and whose logarithm of the octanol-water partition coefficient (logP) is between 1 and 3.

To score the obtained molecules, we trained additional predictive models using two new unsupervised pre-training run. The first unsupervised training was done with the molecules generated in the first six RL runs, the second with the generated molecules from all RL runs. Both were fine-tuned with the CDK9 dataset on the two random splits that were used before, thus, we had four additional predictive models and seven in total with the three models trained earlier.

The molecules were then scored with these seven trained predictive models and the obtained values were used as a basis for selection. The number of molecules that were indicated to be active by at least one model was 4100.

We filtered out all molecules that were not *in-distribution* according to the out-of-distribution classifier. To select compounds to synthesize and test, the remaining molecules were clustered with DataWarrior [36], and we manually selected 69 representatives (listed in Section A.8) from promising clusters.

The selected molecules were purchased from Enamine’s REAL dataset, accessed September 2019, and the activity of the molecules against human CDK9 was assessed using KINOMEscan (Eurofins, [37]).

## A.8 Structures of compounds selected for testing

## A.8.1 Round 1, 207 compounds

~~~
GCCC(CC)c1nnc(NC(=O)N2CCCC[C@H]2c2cn[nH]c2)s1
C[C@@H](CO)N(C)C(=O)NCC1(c2cc(F)cc(C(F)(F)F)c2)CC1
Cc1c(CNC(=O)CCn2cnc3sc4c(c3c2=O)CCCC4)cnn1C
O=C(Cc1csc(-c2ccoc2)n1)Nc1cc(C(=O)O)ccc1F
N[C@@H]1CCN(C(=O)Cn2cnc3scc(-c4cccs4)c3c2=O)C1
N#Cc1cc(COc2cccc(F)c2C(N)=O)ccc1F
CC(C)c1ccc(-c2nc(C(=O)N(C)CC3(CO)CC3)cs2)cc1
Cc1nc(-c2ccc(F)cc2)sc1[C@H](C)NC(=O)N1CCC(C)(O)CC1
Cc1nc(-c2cccc(Cl)c2)sc1C(=O)Nc1ccc(F)cn1
CS(=O)(=O)c1cccc(-c2csc(-c3ccccc3O)n2)c1
CN(Cc1cccnc1)C(=O)Nc1cc(Cl)cc2c1OCC2
COc1ccc(C#N)cc1Cn1cnn(C(C)(C)C)c1=O
CS(=O)(=O)CC1(CNC(=O)N2CC[C@H](Oc3cccc(Cl)c3)C2)CC1
COc1ccc(Cl)cc1NC(=O)Nc1ccn(CCC#N)n1
O=C(NCC1(CO)CCOCC1)c1cc(C2CC2)nc2ccc(F)cc12
CN(C)c1ccc(-c2noc(CC3(CS(C)(=O)=O)CC3)n2)cc1
CCCn1c(=O)n(CCC(=O)N2CCOC(C)(C)C2)c2ccccc21
CN(CC1(CO)CC1)C(=O)c1csc(-c2ccc(Cl)c(Cl)c2)n1
CS(=O)(=O)Nc1cccc2c1CCN(C(=O)Nc1cc(F)cc(Cl)c1)C2
COc1ccc(F)cc1NCc1nc(-c2cnn(C)c2)cs1
CN(C(=O)CNC(=O)c1ccc(F)cc1F)C1(C#N)CCC1
Cc1ccc(Cl)cc1S(=O)(=O)Nc1c(C(N)=O)n[nH]c1C
CCN1CN(C(=O)Nc2ccc3nc(C4CC4)sc3c2)CC1=O
CC1(CNC(=O)c2cc(-c3ccco3)nc3c(C(N)=O)ncn23)CCC1
CS(=O)(=O)Nc1cc(F)ccc1NC(=O)c1cccn1C1CC1
CS(=O)(=O)CC1(Cc2nc(-c3ccc(F)c(F)c3)no2)CC1
Cc1c(NC(=O)Nc2ccn(CCC#N)n2)cccc1N(C)C
COc1cc(F)c(NC(=O)c2cnc(-c3ccccn3)s2)cc1F
CCn1cc(C(=O)N[C@@H](C)CC#N)c(=O)c2cc(Cl)ccc21
Cn1cncc1C1CC(NC(=O)Nc2ccc(F)cn2)CCO1
O=C(CS(=O)(=O)[C@H]1CCc2c(F)cc(F)cc21)Nc1ccccc1Cl
O=C(Nc1cc(Cl)cc2c1OCC2)N1CCC(c2ccncc2)CC1
Cc1ccc(Cl)c(OCC(O)CN2CC(C(N)=O)CCC2C)c1
N#Cc1ccc(NCc2cccnc2N2CCC(C(N)=O)CC2)cc1Cl
CC1(CNc2ccccc2CN2CCC[C@H](C(N)=O)C2)CCOCC1
COc1ccc(Cl)cc1CN(C)C(=O)N[C@H]1CCc2[nH]ncc2C1
C[C@H](NC(=O)Nc1ccc(F)cc1C#N)c1ccccc1NS(C)(=O)=O
CCn1cc(C(=O)N[C@H](C)C(=O)NC)c(=O)c2cc(Cl)ccc21
COc1cc(NC(=O)N(C)Cc2cnn(C)c2)ccc1Cl
CC(=O)Nc1ccc(Cl)c(NCc2nc(-c3cccs3)n[nH]2)c1
CCn1cc(C(=O)NCC(C)(C)C(N)=O)c(=O)c2cc(Cl)ccc21
N#Cc1ccccc1N1CCN(C(=O)Nc2ccc(F)cn2)CC1
CC1CCC(N2C[C@@H](C(=O)Nc3ccc(-c4cnco4)cc3)CC2=O)CC1
COc1ccc(F)cc1C(C)NC(=O)N(C)CC1CCCCC1O
CCn1cc(C(=O)NCC(=O)NC2CC2)c(=O)c2cc(Cl)ccc21
Cc1nc(-c2ccc(F)cc2)sc1[C@H](C)NC(=O)N1CCN(C2CC2)CC1
CC(C)(C)n1ncn(Cc2cc(F)ccc2C#N)c1=O
CC[C@@H](CC#N)NC(=O)Nc1cccc(N(C)C(C)C)c1
O=C(Nc1cccc2c1OCCO2)c1ccc(=O)n(-c2ccccc2F)n1
Cc1cnc([C@@H](C)CNC(=O)NCc2cccc(N(C)C)c2)s1
Cn1nccc1C1CC2CCC(C1)N2C(=O)c1ccoc1
Cc1ccc(S(=O)(=O)Nc2c(C(N)=O)n[nH]c2C)c(Cl)c1
COc1cccc(-c2nc(C(=O)N(C3CC3)[C@H]3CCS(=O)(=O)C3)c(C)[nH]2)c1
NS(=O)(=O)c1cc(C(=O)Nc2ccc(F)cc2OCC2CC2)co1
CCC(CC)n1ncc(C(=O)N[C@H]2COc3cccc(F)c32)c1C
Cc1cc(F)ccc1-c1noc(CC2(CS(C)(=O)=O)CC2)n1
CC(C)N1C[C@@H](C(=O)Nc2ccc(-c3cnco3)cc2)CC1=O
NC(=O)c1ncn2c(C(=O)NCCC3CCCC3)cc(-c3ccco3)nc12
Cc1[nH]nc(C(N)=O)c1NS(=O)(=O)c1cc(F)ccc1Cl
NC(=O)c1ccc(NC(=O)c2cnc(-c3cccs3)s2)c(F)c1
CC(=O)N(C)c1cccc(NC(=O)NC[C@H](C)c2ncc(C)s2)c1
O=C(Nc1cc(Cl)c(O)c(Cl)c1)Nc1ccc(F)cc1F
O=C(CNC1(c2noc(C3CC3)n2)CCCC1)Nc1ccc(F)cc1Cl
COc1cc(F)c(NC(=O)NCC2(CCO)CCCCC2)cc1F
Cc1cnc([C@@H](C)CNC(=O)Nc2cccnn2)s1
COc1ccc(C)cc1CS(=O)(=O)Cc1nc(C2CC2)n[nH]1
Cc1nc2c(s1)CCC[C@@H]2CNC(=O)Nc1ccc(F)cn1
CCn1cc(C(=O)NCC(C)(C)N(C)C)c(=O)c2cc(Cl)ccc21
CN(C)C(=O)COc1ccc(NC(=O)Nc2ccc(F)cc2)cc1
COc1ccc([C@H]2CCN(C(=O)Nc3ccc(F)cn3)C2)cc1
CCn1c(NCc2cc(C(N)=O)cs2)nc2cc(F)ccc21
CCn1cc(C(=O)NCC(C)(C)CO)c(=O)c2cc(Cl)ccc21
Cc1ccnn1-c1ccccc1NC(=O)Nc1ccn(CCC#N)n1
Cc1nc(-c2ccc(F)cc2)sc1[C@H](C)NC(=O)c1ccncc1Cl
O=C(Nc1ccc(F)cc1Cl)c1cnc(-c2cccs2)s1
N#Cc1c(F)cccc1NCc1cccnc1N1CCC(C(N)=O)CC1
CCn1cc([C@@H]2CS(=O)(=O)CCN2C(=O)c2cncc(F)c2)cn1
CC(C)(C)n1ncn(Cc2cc(C#N)ccc2F)c1=O
N#Cc1ccc(NCc2cccnc2N2CCC(C(N)=O)CC2)c(F)c1
CC(C)(C)n1ncn(Cc2ccc(F)c(C#N)c2)c1=O
Cc1ccc(C)c(NC(=O)CN2CCC[C@H](c3nncn3C3CC3)C2)c1
N#CCC(=O)N1CCCC[C@H]1C(=O)Nc1ccc(Cl)cc1Cl
O=C(NCC1(CO)CCC1)Nc1cc(C(F)(F)F)ccc1N1CCCC1
CN(Cc1nc(-c2ccc(F)cc2)no1)C1(CO)CCOCC1
CCn1cc(C2CCN(C(=O)NCCc3csc(C(C)C)n3)CC2)cn1
C[C@H](Nc1cnn(C(C)(C)C)c1)C(=O)Nc1cc(F)ccc1F
CCc1c(NC(=O)[C@H]2CCC(=O)N2)cnn1-c1cccc(Cl)c1
Cc1ccc(-c2ccc(C(=O)NCCn3cnnc3C3CC3)s2)o1
CS(=O)(=O)Nc1ccccc1NC(=O)c1cccn1C1CCCC1
COc1cc(Cl)c(Cl)cc1NC(=O)N1CCC[C@H](C(N)=O)C1
CCn1cc(C(=O)N(C)c2cnn(C)c2)c(=O)c2cc(Cl)ccc21
C[C@H](NC(=O)Nc1cccc2c1OC(C)(C)C2)C1CC1
CCC(CC)n1nc(C(=O)N[C@H]2COc3cccc(F)c32)cc1C
O=C(CNC1(c2noc(C3CC3)n2)CCCC1)Nc1ccc(Cl)cc1
Cc1nc(-c2ccccc2)sc1NC(=O)N[C@@H](C)c1nnnn1C1CC1
Cn1cc(NC(=O)N[C@H]2CC(=O)N(c3ccc(Oc4ccccc4)cc3)C2)cn1
COc1ccc(Cl)c(NC(=O)N(CCO)C(C)C)c1
Cn1nc(C2CCCC2)cc1NC(=O)c1cncc(O)c1
CN(Cc1nc(-c2ccccc2)no1)C1(CO)CCOCC1
C[C@H](Nc1cccc(CN2CCC(C(N)=O)CC2)c1)c1cccnc1
CN(C)C(=O)CCNC(=O)Nc1ccc(Oc2ccccc2C#N)cc1
C[C@H](Nc1cnn(C(C)(C)C)c1)C(=O)Nc1ccc(F)cc1F
NC(=O)C1CCN(C(=O)c2csc(-c3cccc(Cl)c3)n2)CC1
O=C(Nc1ccc(F)cc1)Nc1cc(Cl)c(O)c(Cl)c1
CN(Cc1nc(-c2ccc(Cl)cc2)no1)C1(CO)CCOCC1
N#Cc1cccc(N2C(=O)CN(Cc3ccc(F)cc3)C2=O)c1
Cn1cc(CNc2ccccc2N2CCC(C(N)=O)CC2)cn1
CC(=O)Nc1ccc(Cl)c(NC(=O)Nc2ccn(C)n2)c1
Cc1cnccc1NC(=O)Nc1cc(Cl)c(O)c(Cl)c1
Cc1ccn(CC(=O)N2CC(O)CC2c2cc(F)ccc2F)c(=O)c1
Cn1ccc(NC(=O)N[C@H]2CC(=O)N(c3ccc(Oc4ccccc4)cc3)C2)n1
COc1ccc(NC(=O)c2nn(-c3ccccc3F)cc2O)cc1F
COc1ccccc1CN(C)C(=O)Nc1ccncc1F
c1cc(-c2noc([C@H]3CCn4cncc4C3)n2)cs1
N#Cc1cc(F)ccc1NC(=O)NCc1ccncc1
O=C(NCc1nnc(-c2ccccc2)s1)N1CCC[C@H]1c1cccs1
Cc1nc(-c2ccc(Cl)cc2)sc1C(=O)N[C@@H](C)c1nnnn1C1CC1
Cc1csc(C(C)(C)NC(=O)Cn2c(=O)cnc3ccccc32)n1
CN(C)c1nccc(N2C[C@@H](F)C[C@H]2CNC(=O)NCc2ccco2)n1
NC(=O)[C@H]1CCCN(C(=O)Nc2cc(F)ccc2OCC2CC2)C1
O=C(CNC(=O)Nc1cccnc1-c1ccccc1)NCC1CC1
NC(=O)C1CCN(c2ccccc2NCc2cccnc2)CC1
CN(CCN(C)c1ccccc1)C(=O)c1cnc(-c2ccco2)s1
Cc1c(Cl)cccc1NC(=O)CN1CCC[C@H](c2nncn2C2CC2)C1
Cc1c(C(=O)Nc2ccc(F)cc2C#N)cnn1-c1ccccn1
CN(C)c1nccc(N2C[C@@H](F)C[C@H]2CNC(=O)Nc2ccsc2)n1
C[C@H](NC(=O)Nc1cnn(Cc2ccccn2)c1)c1cccc(O)c1
Cc1csc([C@@H](C)CNC(=O)Nc2ccccc2-n2nccc2C)n1
Cc1nc(-c2ccc(F)cc2)sc1C(=O)N[C@@H](C)c1nnnn1C1CC1
CC(C)n1cnnc1-c1ccccc1NC(=O)Nc1cc(CC(C)(C)C)[nH]n1
CCC(CC)n1nc(C)cc1C(=O)N1CCOc2ccc(F)cc2C1
Cc1cnc(C(C)(C)NC(=O)N[C@H]2CCN(c3ccc(Cl)c(F)c3)C2)s1
O=C(Nc1ccc(O)cc1)Nc1nc2ccc(F)cn2n1
N#Cc1cc(F)ccc1NC(=O)N[C@@H](CO)Cc1ccccc1
O=S(=O)(Nc1nncn1Cc1ccccc1)c1cc(F)ccc1Cl
Cn1nccc1CNc1ccccc1N1CCC(C(N)=O)CC1
Cc1cnccc1NC(=O)NC[C@@H]1C[C@H](F)CN1c1ccnc(N(C)C)n1
CS(=O)(=O)CC1(Cc2nc(-c3ccccc3Cl)no2)CC1
O=c1ccc(F)cn1Cc1ncc(-c2ccccc2F)o1
N#Cc1ccccc1Oc1ccc(NC(=O)NCCCO)cc1
COc1cccc(CNC(=O)Nc2cc(Cl)ccc2-c2nc(C3CC3)no2)n1
Cn1cc([C@H]2OCCC[C@@H]2NC(=O)Nc2ccc(F)cn2)cn1
CCC(CC)n1nc(C)cc1C(=O)N[C@H]1CCOc2ccc(F)cc21
CCc1nc(C2CCN(C(=O)c3ccc(-c4ccco4)[nH]c3=O)CC2)n[nH]1
CCn1cc([C@@H]2CS(=O)(=O)CCN2C(=O)NCCC2=CCCC2)cn1
COc1cccc(F)c1NC(=O)Nc1cnn(C2CCCC2)c1C
CCC(CC)n1nc(C(=O)N[C@H]2CCOc3ccc(F)cc32)cc1C
Cc1n[nH]cc1NC(=O)Nc1ccnn1Cc1ccc(Cl)c(F)c1
CC[C@@H](CC#N)NC(=O)Nc1ccc(NC(C)=O)cc1Cl
C[C@H](NC(=O)Nc1cccc2c1OC(C)(C)C2)c1nnnn1C
O=C(Nc1ccc(F)cn1)N1CCC[C@H]1c1cccs1
O=S(=O)(Cc1nc(C2CC2)n[nH]1)c1nccn1-c1cccc(F)c1
Cc1cc(Cl)c(NC(=O)N[C@@H](C)C(=O)N(C)C)cc1Cl
O=C1N[C@@H](Cc2c[nH]c3cc(F)ccc23)C(=O)N1c1ccccc1Cl
Cc1cc(C)n(-c2ccc(Cl)c(C(=O)Nc3ccc(F)cn3)n2)n1
CN(CCn1cccn1)C(=O)c1cnc(-c2ccco2)s1
COc1cc(F)ccc1NC(=O)c1cn(C)nc1-c1ccccc1Cl
Cc1nnnn1-c1cc(NC(=O)NCCc2ccnn2C)ccc1F
NC(=O)[C@H]1CCCN(C(=O)Nc2cc(F)c(OC(F)F)cc2F)C1
C[C@H](NC(=O)N1CCC(c2ccn[nH]2)CC1)c1c(F)cccc1Cl
CCn1nc(C)c(CNC(=O)N(C)Cc2ccc(OC)c(F)c2)c1C
CC(C)(NC(=O)N[C@H](c1nccs1)C1CC1)c1cn(-c2ccccc2)nn1
O=C(Nc1ccc(F)cn1)N1CCC[C@H](c2nnc3n2CCC3)C1
Cc1ccn(CC(=O)N(C)Cc2ccccc2F)c(=O)c1C#N
COc1cnccc1C(C)NC(=O)Nc1ccnn1C(C)C1CC1
CCn1c(NCc2cnn(C)c2)nc2cc(F)ccc21
CCn1c([C@@H]2CCCN2C(=O)Cc2cccc(O)c2F)nc2ccccc21
O=C(Nc1ccn(Cc2ccncc2)n1)Nc1cc(Cl)ccc1F
NC(=O)c1ccc(NC(=O)N(Cc2c(F)cccc2F)C2CC2)cn1
C[C@H](NC(=O)NC1CCN(C2CC2)CC1)c1ccc(Oc2ccccc2)c(F)c1
CCn1cc([C@H]2OCCC[C@@H]2NC(=O)Nc2ccon2)cn1
CN(Cc1ccc(C#N)cc1)C(=O)Nc1ccncc1F
Cc1nnnn1-c1cc(NC(=O)Nc2ccn(C)n2)ccc1F
Cc1nn(C(C)(C)C)c(=O)n1Cc1nc(-c2ccco2)no1
Cc1c(C(=O)N[C@@H](CS(C)(=O)=O)c2ccccc2)cnn1C1CCCC1
Cc1nnc(Cn2cnc3c(cnn3C(C)(C)C)c2=O)n1C1CC1
Fc1ccc(-c2noc([C@H]3CCn4ccnc4C3)n2)c(Cl)c1
NC(=O)[C@H]1CCCN(C(=O)Nc2ccc(-c3ccco3)cc2F)C1
Cc1c(C(=O)N2c3ccccc3OC[C@@H]2C)nnn1Cc1ccc(F)cc1F
Cc1c(C(=O)N2c3ccccc3C[C@H]2C(N)=O)nnn1Cc1ccc(F)cc1F
Cc1c(C(=O)N2CC(C)(C)Oc3ccccc32)nnn1Cc1ccc(F)cc1F
C[C@H](NC(=O)c1csc(-c2ccccc2Cl)n1)c1nnnn1C1CC1
CCn1cc(NC(=O)Nc2ccnc(C)c2Cl)cn1
O=C(Nc1cccnn1)N[C@@H](Cc1ccccc1)C1CC1
NS(=O)(=O)Cc1ccc(NC(=O)c2cccn2C2CCCC2)cc1
CN(C)c1ncc(NC(=O)NC(C)(C)c2cn(-c3ccccc3)nn2)cn1
C[C@@H](NC(=O)Nc1ccc(F)cn1)c1ccccc1
CCc1nc2cc(CNC(=O)c3ccco3)ccc2n1C1CC1
O=C(Nc1ccccn1)N[C@@H](Cc1ccccc1)C1CC1
O=C(CCn1cnc2ccccc2c1=O)N(c1ccccn1)C1CCCC1
CCc1ncc(NC(=O)NC(C)(C)c2cn(-c3ccccc3)nn2)cn1
CCS(=O)(=O)C[C@H](C)NC(=O)C(C)(C)c1ccc(Cl)cc1F
C[C@H](NC(=O)N(C)CC1(O)CCOCC1)c1ccc(Cl)s1
CCS(=O)(=O)c1cc(F)ccc1NC(=O)NCc1ccco1
C=C(C)CNC(=O)Nc1cccc(CS(C)(=O)=O)c1
Cc1nn(C(C)(C)C)c(=O)n1CCOc1ccccc1Cl
CNC1CCN(C(=O)c2nn(-c3ccc(F)c(F)c3)c3c2CCC3)CC1
CN1CCC(NC(=O)Nc2cnn(C(C)(C)C)c2)C1c1ccccc1
C[C@H](Oc1ccc(C(C)(C)C)cc1)C(=O)N1CCC(C(N)=O)CC1
CC(C)(NC(=O)Nc1ccncc1F)c1cn(-c2ccccc2)nn1
NC(=O)[C@H]1CCCN(C(=O)Nc2ccc(F)cc2F)C1
NC(=O)c1ncn([C@H]2CCCN(C(=O)NCc3ccoc3)C2)n1
O=C(c1n[nH]c2c1CCC2)N(Cc1ccsc1)[C@H]1CC12CCNCC2
O=C(NCC(=O)N1CCCC1)Nc1ccccc1Oc1ccccc1
CN(C)C[C@H](NC(=O)NCc1ccoc1)c1ccc(Cl)cc1
Cc1c(C(=O)N(C)Cc2ccc(F)c(F)c2)nnn1C1CCNCC1
Cc1c(C(=O)NCCc2ccc(F)cc2F)nnn1C1CCNCC1
~~~

## A.8.2 Round 2, 69 compounds

~~~
CC1=C(C2=CC=CC=C2)SC(NC(=O)C2CCCCNC2)=N1
COC1=CC=CC=C1C1=CC=NC(NC[C@H]2C[C@@H]3CC[C@H](C2)N3)=N1
O=C(CC1=CC(F)=CC=C1)NCC(=O)N1CC=C(C2=C[NH]C3=NC=CC=C23)CC1
O=C(N1CC=C(C2=C[NH]C3=NC=CC=C23)CC1)N1CCC1
COC1=CC=C(C2=CC=NC(NC(=O)C3=NC=C(Cl)C=N3)=N2)C=C1
O=C(CNC1=CC=CC=N1)N1CC=C(C2=C[NH]C3=NC=CC=C23)CC1
CC(C)C(O)CC(=O)N1CC=C(C2=C[NH]C3=NC=CC=C23)CC1
COC1=CC=C(C)N=C1NC(=O)CC1=NOC2=CC=C(Br)C=C12
COC1=CC=C2ON=C(CC(=O)NC3=NC(C)=CC=C3O)C2=C1
CCC(CNC(=O)CC1=NOC2=CC=CC=C12)NC(=O)CC1=NOC2=CC=CC=C12
CC1=CON=C1NC(=O)CC1=NOC2=CC=C(Br)C=C12
NC(=O)C1(CNC2=NC=NC3=CC=CC(Br)=C23)CCOCC1
NC(=O)C1(CNC2=NC=NC3=C2C2=CC=CC=C2[NH]3)CCOCC1
COC1=CC=CC=C1C1=CC=NC(NC(=O)C2=CC=C(NS(C)(=O)=O)N=C2)=N1
CC(C)COCC(O)CN1C=NC2=C(NCC3=CC=CC=C3)N=CN=C21
O=C(COC1=CC=CC=C1C1=CC=CC=C1)NC1=CN=CC=N1
O=C(COC1=CC=CC=C1C1=CC=CC=C1)NC1=NC=CC=N1
O=C(COC1=CC(F)=CC(F)=C1)NC1=NN=C(C2=CC=CC=C2Cl)[NH]1
COC1=CC=CC=C1C1=NN=C(NC(=O)COC2=CC(F)=CC(F)=C2)[NH]1
O=C(COC1=CC=C(F)C=C1Cl)NC1=CC=CC=C1N1N=CC2=C1N=C[NH]C2=O
O=C(COC1=CC=C(F)C=C1Br)NC1=CC=C(N2N=CC3=C2N=C[NH]C3=O)C=C1
COC1=CC=C(Br)C=C1SCCCN1C=NC2=C(N)N=CN=C21
N#CCCN1N=CC2=C(NC3=CC=C4NC(=O)CC4=C3)N=CN=C21
O=C(CC1=C[NH]C2=NC=CC=C12)NC1=NOC(C2=CC=CC=C2)=C1
ClC1=CC=CC(CN2N=CC3=C(NCCNC4=CN=CC=N4)N=CN=C32)=C1
COC1=CC=C(Cl)C=C1NC(=O)CCN1C=NC2=C(N)N=CN=C21
C[C@H]1CN(C(=O)N2CC=C(C3=C[NH]C4=NC=CC=C34)CC2)C[C@@H](C)O1
NC(=O)C1(CNC2=NC=NC3=C2C=NN3C2=CC=CC(Br)=C2)CCOCC1
O=C(COC1=CC(F)=CC(F)=C1)NC1=NN(C2=CC=NC=C2)C=C1
CC1=N[NH]C2=NC=C(NC(=O)CC3=NOC4=CC=CC=C34)C=C12
O=C(COC1=CC(F)=CC(F)=C1)NC1=NC2=CC=CN=C2S1
O=C(CC1=NOC2=CC=CC=C12)NC1=CC=CC(C2=C[NH]N=C2)=C1
CC1=NC(NCC2=CC=CC(OCC(N)=O)=C2)=C2C=NN(C3=CC=CC(F)=C3)C2=N1
NC1=NC=NC2=C1C=NN2CC1=CSC(C2=CC=C(F)C=C2)=N1
COC1=CC=C(C2=NN=C(NC(=O)CC3=NOC4=CC=CC=C34)[NH]2)C(OC)=C1
COC1=CC=CC2=C1OC(CNC1=C3C=NN(CCC#N)C3=NC=N1)=C2
CN1N=CC2=C1N=C(N[C@H]1C[C@@H](NC(=O)CC3=NOC4=CC=CC=C34)C1)[NH]C2=O
O=S(=O)(NC1=NC2=CC=CC=C2N=C1NCCC1CC(O)C1)C1=CC=CC=C1
CN(C)CC1=CC=NC(NC(=O)COC2=CC=C(F)C=C2Cl)=C1
CCN(CC1=CC=C2OCCOC2=C1)C(=O)CN1C=NC2=C(N)N=CN=C21
O=C(COC1=CC=C(Br)C=C1F)NC1=NC=C(Cl)C(O)=C1
COC1=CC(F)=CC=C1C1=CC=C(NC(=O)C2CCCNC2)N=C1
CC1=CC(NC(=O)COC2=CC=C(Br)C(F)=C2)=NO1
NC1=NC=NC2=C1N=CN2CCCOC1=NN=C(C2=CC=CC=C2)O1
O=C(COC1=CC=CC=C1F)NC1=CC=C(N2C=NC=N2)C=N1
O=C(COC1=CC=CC(Br)=C1)NC1=N[NH]C(CC2=CC(F)=CC=C2)=N1
CC1=CC=NC(NC(=O)CCC2=C(C)C3=C([NH]C2=O)N(C)N=C3C)=C1
COC1=CC=C(OCC(=O)NC2=NN=C(C3=CC=CC=C3OC)[NH]2)C=C
CNC1=NN=C(SCC2=NC3=CC=CC=C3C(=O)[NH]2)S1
CC(NC(=O)CCN1C=NC2=C(N)N=CN=C21)C1=CC=CC=C1
C[C@H](NC(=O)CCN1C=NC2=C(N)N=CN=C21)C1=CC=CC=C1
O=C(CC1=C[NH]C2=NC=CC=C12)NC1=CC=C(NC2=NC=CC=N2)C=C1
CNC1=NN=C(SCC2=NC(N)=C3C=CC=CC3=N2)S1
COC1=CN=C(NS(=O)(=O)C2=CC=C(NC(=O)CC3=NOC4=CC=CC=C34)C=C2)N=C1
COC1=CC=C2ON=C(CC(=O)NC3=NN=C4C=C(C)C=CN34)C2=C1
COC1=CC=CC=C1C1=CC=NC(NC(=O)COC2=CC=CC=C2)=N1
COC1=CC=CC=C1OCC(=O)NC1=CC=C(N2N=CC3=C2N=C[NH]C3=O)C=C1
O=C(COC1=CC=CC=C1F)NC1=CC=C2OCC(=O)NC2=N1
COC1=CC=C(Cl)C=C1NC(=O)COC1=NC=NC2=C1C=NN2C1=CC=CC=C1
CC1=NOC2=NC=C(NC(=O)C3=CC=C(NC(=O)CC4=CC=CN=C4)C=C3)C=C12
O=C(CC1=CC=CC(OCC2=CC=CN=C2)=C1)NC1=CC=C(S(=O)(=O)NC2=NC=CC=N2)C=C1
COC1=CC(OC)=CC(C2=CC=CC(NC(=O)C3CCCOC3)=N2)=C1
CCOC1=CC=CC=C1OCC(=O)NC1=CC=CC(OCC(F)F)=N1
O=C(COC1=CC=C(Br)C=C1F)NC1=CN=CC=N1
CC1=CC(C)=NC(NC(=O)CCC2=CC=C3OCC(=O)NC3=C2)=C1
O=C(COC1=CC(Cl)=CC=C1)NC1=NN(CC2=CC=NC=C2)C=C1
N#CCCN1N=CC2=C(NCC3(C4=CC=C(F)C=C4)CC3)N=CN=C21
NC(=O)C(CNC1=NC=NC2=C1C=NN2C1=CC=NC=C1)CC1=CC=CC=C1Cl
COC1=CC=CC=C1CCNC(=O)CCN1C=NC2=C(N)N=CN=C21
~~~

## Footnotes

1 30 *μ*mol is equivalent to 30 micromoles of the compound in 1 liter of solution for the assay.

2 The fact that the Jensen-Shannon mutual information is essentially a cross-entropy loss function makes training more stable.

